# Design of linked-domain protein inhibitors of UBE2D as tools to study cellular ubiquitination

**DOI:** 10.1101/2024.09.02.610852

**Authors:** Zara Bukhari, Li Gu, Anneroos E. Nederstigt, Logan J. Cope, Derek L. Bolhuis, Kim Harvey, Tristan Allen, Spencer Hill, Yujie Yang, Guy Lawson, Cai Lu, Tommy Tran, Leah Pineda, Leanne Low, Andrew Chiang, Jason Song, Michelle V. Fong, Vanessa M. Rangel, William K. Chan, Gary Kleiger, Dennis Goldfarb, Craig A. Vierra, Nicholas G. Brown, Joseph S. Harrison

## Abstract

Ubiquitin (Ub) is a post-translational modification that largely controls proteostasis through mechanisms spanning transcription, translation, and notably, protein degradation. Ub conjugation occurs through a hierarchical cascade of three enzyme classes (E1, E2, and E3s) involving >1000 proteins that regulate the ubiquitination of proteins. The E2 Ub-conjugating enzymes are the midpoint, yet their cellular roles remain under-characterized, partly due to a lack of inhibitors. For example, the cellular roles of the promiscuous E2 UBE2D/UBCH5 are not well described. Here, we develop a highly selective, multivalent, engineered protein inhibitor for the UBE2D family that simultaneously targets the RING- and backside-binding sites. In HeLa cells, these inhibitors phenocopy knockdown of UBE2D by reducing the IC_50_ to cisplatin and whole-cell proteomics reveal an increased abundance of ∼20% of the identified proteins, consistent with reduced Ub degradation and proteotoxic stress. These precision tools will enable new studies probing UBE2D’s central role in proteome management.

## Introduction

Ubiquitin (Ub) regulates protein homeostasis, which includes the synthesis, folding, conformational maintenance, assembly, trafficking, function, and degradation of proteins^1–3^. As such, ubiquitination regulates nearly all biological pathways and is dysregulated in many diseases, including multiple hallmarks of cancer^4–6^. Ub conjugation occurs through an enzymatic cascade that begins with an E1-activating enzyme, forming a cysteine-linked thioester intermediate E1∼Ub (∼ denotes a thioester)^7,8^. Next, the Ub conjugate is transferred to one of >40 E2 Ub-conjugating enzymes^9,10^. E2∼Ub collaborate with E3 ligases, of which there are well over 600^11–13^, to ubiquitinate target proteins. Most commonly, Ub is attached to lysines, although other chemical groups have been identified that can be ubiquitinated^14,15^. Given the complexity of the conjugation pathway and the diverse signaling effects of ubiquitination, precisely determining the cellular roles of ubiquitinating enzymes and accessory proteins has been challenging, and new tools are needed.

The E2 Ub-conjugating enzyme family UBE2D is a prime example of the difficulties in studying ubiquitination in the cell. The family consists of four isoforms of UBE2D with >90% sequence similarity, and *in vitro* UBE2D works with virtually all E3 ligases and has been extensively biochemically characterized. However, defining the cellular functions has been more difficult. Since the four isoforms of UBE2D are dispersed on different chromosomes (Ube2D1:10q21.1, Ube2D2:5q31.2, Ube2D3:4q24, and Ube2D4:7p13), genetic knockdown studies are complicated. Also, few small molecule inhibitors are available, none bind to UBE2D with high-affinity (<∼10^-5^ M K_d_), and many do not enter the cell^16–20^. Therefore, it is necessary to develop new tools targeting UBE2D to enable cellular studies to dissect Ub signaling networks.

Here, we unveil a strategy to make potent high-affinity inhibitors for the E2s by mimicking the multivalent binding of E3s. We designed chimeric, domain-linked fusion proteins that consist of a RING/UBOX domain and a ubiquitin-like (UBL) domain, allowing the molecule to bind two sites on UBE2D simultaneously. These proteins have affinities that span 3×10^-6^M-∼1×10^-9^M. Transfecting them into cells reveals significant changes to the proteome, ∼20% of the identified proteins were found to be more abundant compared to ∼3% that were less abundant, which is consistent with reduction of Ub-mediated protein degradation. Gene enrichment analysis of the proteome changes resembles profiles of cells experiencing proteotoxic stress, either from treatment with proteasome inhibitors or protein aggregation diseases like Parkinson’s and Alzheimer’s. We also observed enrichment of many multiprotein complexes and pathways outside of proteasomal degradation, like RNA processing, ribosomal proteins, and non-proteasomal quality control pathways. Cells treated with the inhibitor also have a six-fold reduction in cisplatin IC_50_, demonstrating reduced stress tolerance associated with knocking down/inhibiting UBE2D. Our studies highlight the varied roles of UBE2D and the linked-domain inhibitors described here will enable future studies to dissect the cellular roles of UBE2D in proteostasis.

## Results

### Designing linked domain inhibitors of UBE2D

E2s have a similar fold and core sequence, therefore, we looked to interactions between E2s and E3s to inform our inhibitor design strategy. E3s use multivalent engagement to bind E2s^21–23^. For UBE2D, this occurs by simultaneously recognizing the RING-binding site and a β-sheet surface on the opposite face, called the backside, that was first identified as a weak Ub-binding site^24^, but now several other domains have been found to access this location^22^. One example is the RING E3 ligase UHRF1, which also has a ubiquitin-like domain (UBL) that binds to the backside with ∼20-fold higher affinity than Ub and regulates the ubiquitination of histone H3^25,26^. Biochemical assays show that UHRF1 H3 ubiquitination is specific for UBE2D due to its interaction with UHRF1 UBL. Since the E1s and E3s often recognize UBE2D by engaging multiple sites, we envisioned designing a molecular clamp formed by linking the two binding domains.

We first tested if the isolated UHRF1 UBL and UHRF1 RING domains could inhibit ubiquitination. At high concentrations, the RING (75μM) and UBL (50μM) domains could partially block H3 peptide ubiquitination, and when added together, synergistic inhibition was observed (**Figure S1A and S1B**). Therefore, we proceeded to make the linked domain designs. We estimated that a 4xGGSS linker (∼2.4Å per residue) would be sufficient to span the 34Å between the C-terminus of the RING and N-terminus of the UBL based on a computational model from our previous work **(Figure 1A)**^25^.

**Figure 1:**
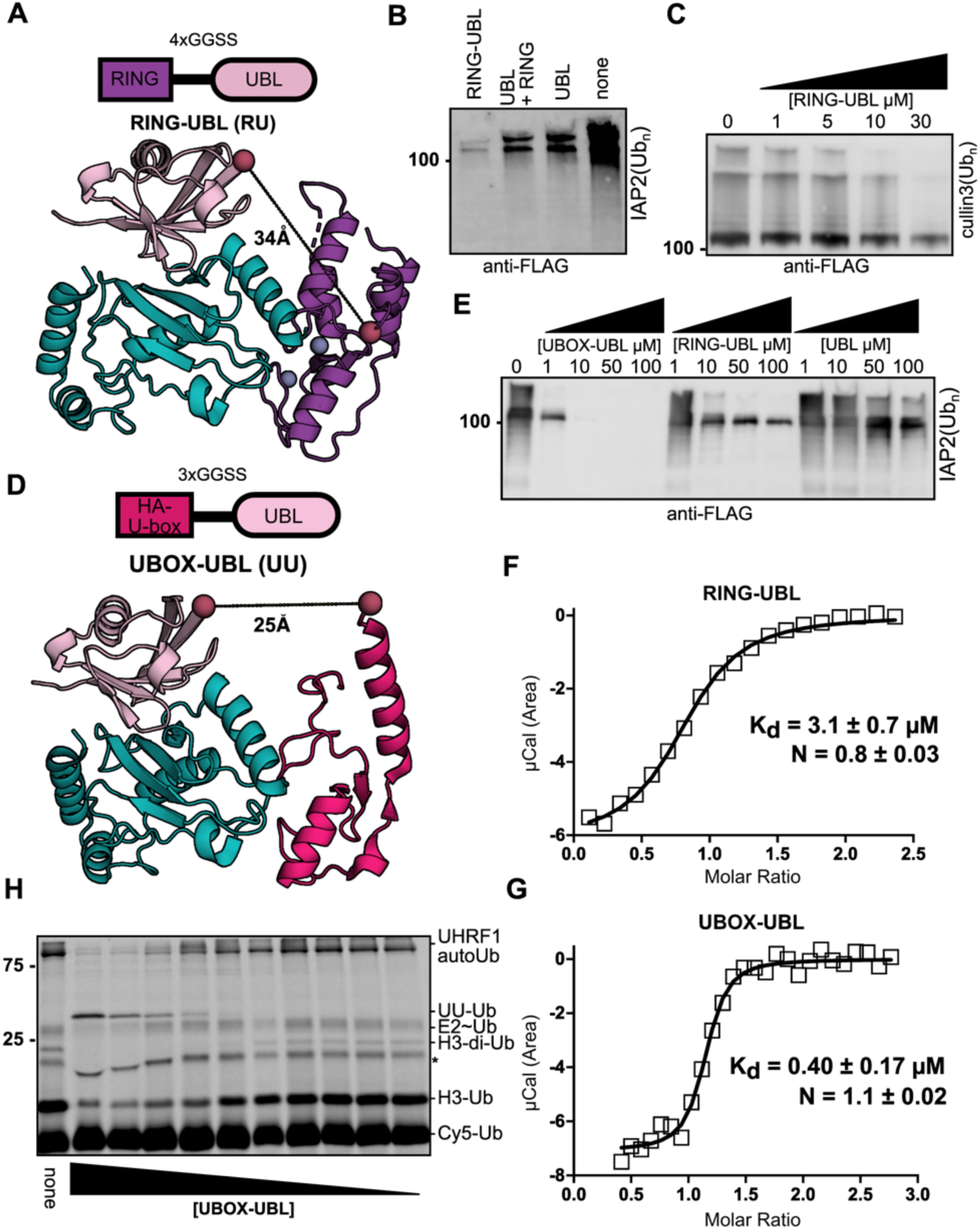
Linked-domain proteins inhibit UBE2D. *A)* Schematic for the design of RING-UBL. The structural model of the UHRF1 RING (purple) bound to UBE2D (cyan) was produced by aligning the UHRF1 RING to the RNF4 RING domain in PDB 4AP4 and the UBL/UBE2D model was produced using Rosetta in a previous publication^25^. The expected distance between the C-terminus of the UHRF1 RING and the N-terminus of the UHRF1 UBL is shown. *B)* IAP2 autoubiquitination assay in the presence of the UBL (50μM), UBL (50μM) and RING (75μM), and RING-UBL (50μM). *C)* Cul3 autoubiquitination in the presence of the indicated amounts of RING-UBL. *D)* Schematic for the design of UBOX-UBL. The structural model of the UHRF1 UBL and E4B UBOX (hot pink) domains bound to UBE2D was produced by aligning the UBOX domain from PDB 2KRE to the RING domain of UHRF1 from our previous model. The expected distance between the C-terminus of the UBOX and the N-terminus to the UHRF1 UBL is shown. *E)* Inhibition of IAP2 autoubiquitination in the presence of the indicated concentrations of UBOX-UBL, RING-UBL, and the UBL domain. Ub assays in B-D were conducted using FLAG-Ub and visualized using anti-FLAG WB. ITC binding isotherm for (F) RING-UBL and G) UBOX-UBL binding to UBE2D1. Thermodynamic parameters and heat per injection shown in Figures S1E-G. H) UHRF1 ubiquitination assay in the presence of the indicated concentrations of UBOX-UBL (90, 45, 15.6, 3.9, 0.975, 0.244, 0.131, 0.087, 0.058, 0.038 μM). This Ub assay was conducted using Cy5-Ub and the * represents a background band in the Ub stock.

We then tested the RING-UBL protein in a variety of E3 (IAP2, UHRF1, CUL3/RBX1) ubiquitination assays, and all show it is more potent than the isolated domains alone or in combination **(Figure 1B, S1C, and S1D)**. For example, in the CUL3/RBX1 autoubiquitination assay, the UBL and RING together have little impact on the reaction, but the RING-UBL can markedly reduce the ubiquitination, indicating that the linking strategy improves the potency of the domains **(Figure S1D)**. We also measured dose-dependent inhibition of RING-UBL, but concentrations of >30μM are needed to observe substantial inhibition, and at concentrations below 10μM, little inhibition is observed **(Figure 1C)**. A more potent inhibitor is needed to be useful inside the cell.

To create a higher affinity inhibitor, we replaced the UHRF1 RING domain, which has a relatively weak affinity for UBE2D (K_d_ ∼75μM)^25^, with a variant of the UBE4B UBOX domain containing two affinity-enhancing mutations identified using phage display^27^. The UBOX has an extended α helix compared to the RING, positioning the C-terminus closer to the N-terminus of the UBL domain, and only requires a 3xGGSS linker to connect the UBOX and UBL domains **(Figure 1D)**. We directly compared RING-UBL and UBOX-UBL in the IAP2 autoubiquitination assay, revealing that UBOX-UBL was a significantly more potent inhibitor, yielding substantial inhibition even at concentrations as low as 1μM **(Figure 1E)**. Next, we used Isothermal Titration Calorimetry (ITC) to determine the affinity of the RING-UBL and UBOX-UBL for UBE2D1. As expected, the UBOX-UBL had a roughly 10-fold higher affinity than the RING-UBL (400nM ± 170nM vs 3.1 ± 0.7μM; **Figures 1F, 1G, and S1E-G**). The RING-UBL had the same affinity as UHRF1 with UBE2D1^28^. Importantly, the N-value in both experiments was near one, indicating 1:1 binding of the inhibitor to UBE2D. We also tested UBOX-UBL in UHRF1 ubiquitination assays using a non-reducing SDS-page gel so we could also observe the UBE2D∼Ub conjugate (**Figure 1H**). In this assay, high concentrations of UBOX-UBL can block UHRF1 autoubiquitination, H3-Ub, and UBE2D∼Ub, suggesting that UBOX-UBL may interfere with Ub transfer from the E1. Additionally, we observe the formation of UBOX-UBL-Ub at higher concentrations.

### Mechanisms of inhibition for the linked-domain UBE2D inhibitors

We used a variety of biochemical assays to further characterize the inhibition mechanism of the linked-domain inhibitors. First, we wanted to confirm that both domains are required for inhibition and tested a known loss-of-function mutation to the UBL domain, F46V^25^ (**Figure 2A; top**). While this UBL mutation did reduce the inhibition, at higher concentrations, we observed only partial inhibition of IAP2 autoubiquitination (**Figure S2A and SB**), especially for the UBOX-UBL variant. To identify a loss-of-function mutation to the RING and UBOX domains, we analyzed the COSMIC (Catalogue of Somatic Mutations in Cancer) database^29^. We found structurally synonymous mutations on a loop in the RING/UBOX at the E2 interface that we anticipate could disrupt binding (**UBOX; Figure 2A, RING; Figure 2B, and Supplemental Table 1**). We tested the RING/UBOX mutations in combination with the F46V-UBL mutation. Both UHRF1 RING substitutions (**Q728H and I725T; Figures S2C and S2D**) and one of the UBOX-UBL (**P1235T; Figure 2E and F**) variants could no longer inhibit ubiquitination assays even at very high concentrations. These mutational experiments demonstrate both domains are necessary for potent inhibition, and these null constructs will be important controls in future cellular studies.

**Figure 2:**
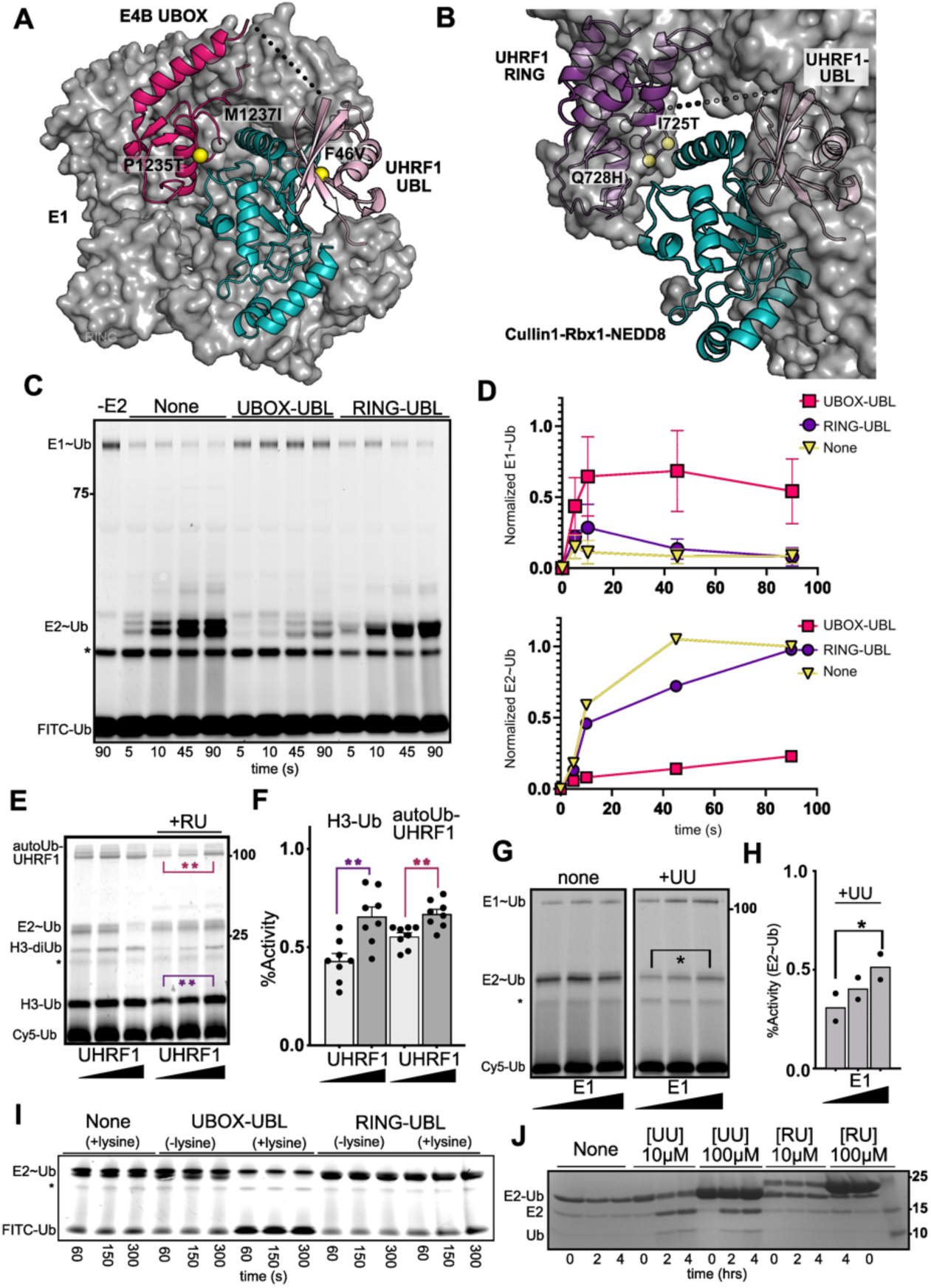
Inhibition mechanism for the linked-domain inhibitors. *A)* Crystal structure of E1 (surface; grey)/Ube2D (cyan) complex (PDB code: 4II2) with the E4B UBOX and UBL model superimposed to show the overlap between the UBOX-UBL and the E1. The tested COSMIC mutations in the UBOX and the F46V mutation in the UBL are shown as spheres. Mutations that abrogated inhibition are shown in yellow. The supporting experiments are shown in Figures S2A, S2C, and S2D. *B)* Crystal structure of the Cul1^nedd8^-Rbx1 (CRL) bound (surface; grey) to UBE2D with the UHRF1 UBL and UHRF1 RING superimposed to show the overlap with the Cul1^nedd8^-Rbx1 (CRL). The Ub conjugate is omitted from the CRL surface map for clarity. COSMIC mutations tested in the UHRF1 RING are shown as spheres. Ubiquitination assays are shown in Figure S2B, S2E, and S2F. *C)* E2 loading assay in the presence of UBOX-UBL and RING-UBL. *D)* Quantification of the E1∼Ub (top; n=2) and E2∼Ub (bottom) in the assay depicted in panel C. *E)* E3 competition assay with increasing UHRF1 concentration (0.7, 2, 4μM) in the presence and absence of 15 μM RING-UBL. *F)* Quantification of normalized H3-Ub and UHRF1 autoubiquitination activity (+inhibitor/-inhibitor) from 0.7μM versus 4μM UHRF1. Statistical significance tested using the repeated-measure one-way ANOVA (**=p-value < 0.01 n=8) *G)* E1 competition assay with increasing E1 concentration (100nM, 200nM, 400nM) in the presence and absence of 1μM UBOX-UBL. *H)* Quantification of normalized E2∼Ub band (+inhibitor/-inhibitor) from the assay depicted in panel G. Statistical significance tested using a repeated-measure one-way ANOVA (*=p-value <0.05 n=2). *I)* EDTA-quenched thioester ubiquitin discharge assay in the presence of 23μM UBOX-UBL and RING-UBL. *J)* Oxyester discharge assay in the presence of UBOX-UBL and RING-UBL at the indicated concentrations. In this assay the bands are detected using Coomassie stain.

Since the *in vitro* ubiquitination assays are multistep reactions, we explored which steps in the reaction could be blocked by the inhibitors. Co-crystal structures have shown that the E1/E2 (**Figure 2A**)^7^ and E3/E2 binding surfaces^30^ (**Figure 2B**) have some overlap, and we anticipated that the linked-domain proteins could interfere with both reactions. To study the E1 transthiolation of E2, we used the UBE2D-loading assay, which contains only E1, UBE2D, Ub, and Mg-ATP, and we monitored the amount of E1∼Ub and E2∼Ub formed. Inhibition of UBE2D loading will lead to the accumulation of E1∼Ub and reduced E2∼Ub, and we observe both of those trends with RING-UBL and UBOX-UBL (**Figures 2C and D**), with UBOX-UBL providing more potent and longer lasting inhibition of E2 loading. On the other hand, the isolated UBL, RING, or UBOX domains did not significantly impact E2 loading (**Figure S2G**).

Next, we tested whether the inhibitors were competitive with respect to E1 and E3. We increased the concentration of E3 (UHRF1) in the presence of RING-UBL (**Figure 2E and 2F**) or E1 in the presence of UBOX-UBL (**Figure 2G and 2H**) to partially overcome the inhibition. In both cases, increasing the concentration of the E3 or E1 decreased the inhibition, suggesting that these inhibitors are competitive with both the E1 and the E3.

Another potential activity of the linked-domain proteins is that they may promote the non-productive discharge of the conjugated Ub by stabilizing the active “closed” conformation of E2∼Ub^31–33^. We used two forms of the UBE2D-Ub conjugate: either thioester, formed from quenching the E2 loading reaction with EDTA, (**Figure 2I**) or purified oxyester conjugate (UBE2D_C85S_-Ub) (**Figure 2J**). We then added free lysine and monitored the reduction of Ub conjugate in the presence or absence of the inhibitors. In these assays, or when using UbΔGG as a substrate (**Figure S2H**), we saw that the UBOX-UBL, but not RING-UBL, enhanced Ub discharge, like the activity of the RING/UBOX domains alone (**Figure S2I**). These results are consistent with our previous work that shows the UHRF1 RING does not accelerate Ub discharge^25,28^.

Therefore, we have shown that the linked-domain inhibitors can interfere with E2 activity in three ways: by preventing the charging of UBE2D, blocking interactions with E3s, and by enhancing non-productive discharge of the Ub conjugate. Due to the multivariate nature of the inhibition, we expect the linked-domain inhibitors to be highly effective in cells.

### RING-UBL and UBOX-UBL are specific for UBE2D

Our previous studies showed that the UHRF1 UBL domain is selective for UBE2D^25^. To determine the selectivity of the linked-domain inhibitors, we began by using substrate ubiquitination assays with two multi-subunit E3s that are capable of using different E2s: Cul1^Nedd8/^Rbx1 mediated β-TrCP ubiquitination of a β-catenin peptide with UBE2D or UBE2R (**Figure 3A and B**)^34^ and APC/C^CDC20^ ubiquitination of Cyclin-B-Ub (**Figure 3C, D and Figure S3A**) using UBE2D, UBE2C, or UBE2S^35,36^. In both substrate ubiquitination assays, the linked-domain inhibitors selectively block UBE2D isoforms and not the other E2s. These assays also demonstrate that the inhibitors can be effective against many E3s, including multisubunit E3s like APC/C and Cullins, which are responsible for ubiquitinating a large portion of the proteome. We also tested SUMO-UBOX autoubiquitination with UBE2D and UBE2E^37^ because the catalytic domains of UBE2D and UBE2E have high amino acid similarity^38^. We only observed inhibition of UBE2D and not UBE2E (**Figure 3E and F**), further underscoring the specificity of the linked-domain inhibitors.

**Figure 3:**
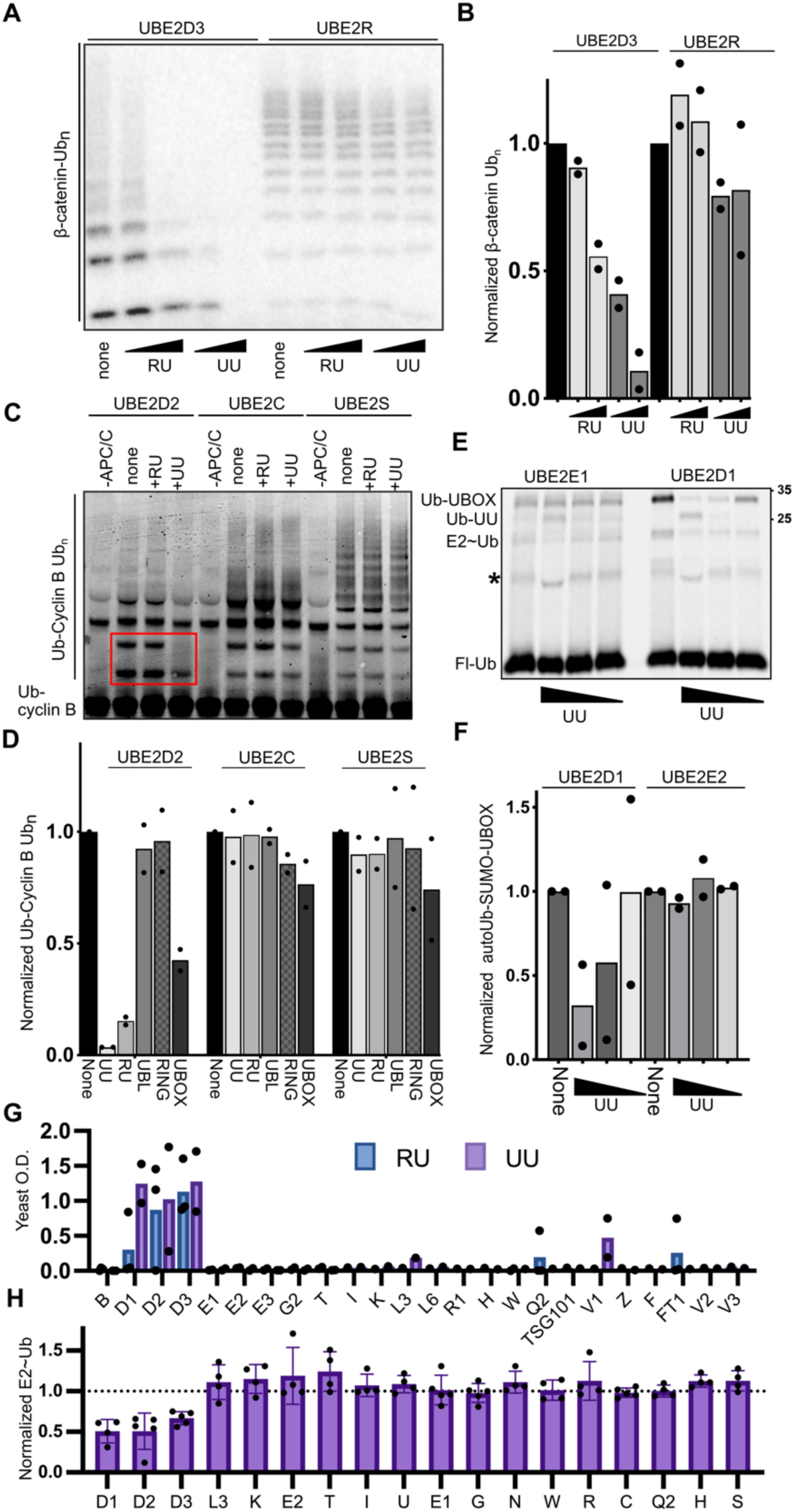
Linked-domain inhibitors are selective for UBE2D. *A)* Skp1/CUL1^Nedd8^/F-box/Rbx1 (SCF) ubiquitination of P32 β-catenin peptide with either UBE2D3 or UBE2R in the presence of 10μM or 100μM of RING-UBL or UBOX-UBL. *B)* Quantification of the ubiquitinated products in A (n=2). *C)* Example APC/C ubiquitylation assay of fluorescent Ub-Cyclin B with either UBE2D2, UBE2C, or UBE2S in the presence of either RING-UBL or UBOX-UBL. *D)* Quantification of APC/C reaction in Figure S3A with 23μM of each inhibitor (n=2). *E)* Autoubiquitination of SUMO-UBOX using UBE2D1 or UBE2E1. UBOX-UBL concentrations are 1, 10, and 100 μM. *F)* Quantification of the assays depicted in panel E (n=2). *G)* Yeast two-hybrid assay showing growth of yeast co-transformed with the inhibitor and a single E2 from the panel of 24, grown in liquid synthetic dropout media lacking Histidine, Tryptophan, and Leucine and supplemented with Aerobasidin A (n=2). *H)* E2 loading assay with the indicated recombinant purified E2s (n=3-5 depending on the E2). While there are no significant differences between D1, D2, D3, all other E2s were significantly different from D1, D2, and most from D3 (p-value <0.05). Statistics are tested using a repeated-measure, one-way ANOVA.

To further probe these inhibitors’ selectivity, we used an established yeast two-hybrid assay that contained 24 E2 proteins^39^. We developed a liquid culture-based growth assay where the yeasts were sequentially transformed with the bait and prey vectors (GAL4AD-E2 and GAL4DNA-BD-linked-domain) and inoculated in the selective condition lacking Histidine and containing Aerobasidin A, in addition to -Leu/-Trp for vector maintenance. This assay had high stringency, and most of the E2 inhibitor combinations did not support yeast growth even after weeks of incubation, and after 7-10 days we measured the optical density (**Figure 3G**). For UBE2D isoforms with the inhibitors we typically observed visible growth within 3-5 days, although for UBOX-UBL, we did observe some growth with the nonfunctional E2 UBE2V1, an E2 adaptor for UBE2N. To directly test for inhibition, rather than only interactions, we performed the E2-loading assays with 18 E2 ubiquitin-conjugating enzymes that we purified in-house. In this assay, we see that UBOX-UBL can inhibit the loading of only the D isoforms, confirming that the linked-domain inhibitors are highly selective (**Figure 3H**).

### Developing nanomolar linked-domain inhibitors using engineered Ub variants

We next sought to develop an even higher affinity inhibitor to UBE2D. Recently, a Ub variant (UbvD1) was selected using phage display that has an affinity of 65nM to the backside of UBE2D1^40^, which is >100 times higher affinity than the UBL domain from UHRF1. We constructed a new linked-domain inhibitor, UBOX-UbvD1_short_, which contained the same 3xGGSS linker used in UBOX-UBL (**Figure 4A**). We tested UBOX-UbvD1_short_ in IAP2 ubiquitination assays and found this new design performs significantly better than the UBOX-UBL or UbvD1 alone, and even at 1μM it almost completely inhibited IAP2 autoubiquitination (**Figure 4B**). On the other hand, the single domain of UbvD1 did reduce polyubiquitination, but could not completely inhibit IAP2 autoubiquitination at any concentration, whereas at 10μM UBOX-UBL almost completely inhibits autoubiquitination, despite having approximately 10-fold weaker affinity (**Figure 4C**). In UHRF1 assays, UbvD1 could only inhibit H3 peptide ubiquitination and not UHRF1 autoubiquitination (**Figure S4A**) further indicating a limitation of solely backside binding inhibitors compared to the multivalent inhibitors like UBOX-UbvD1_short_.

**Figure 4:**
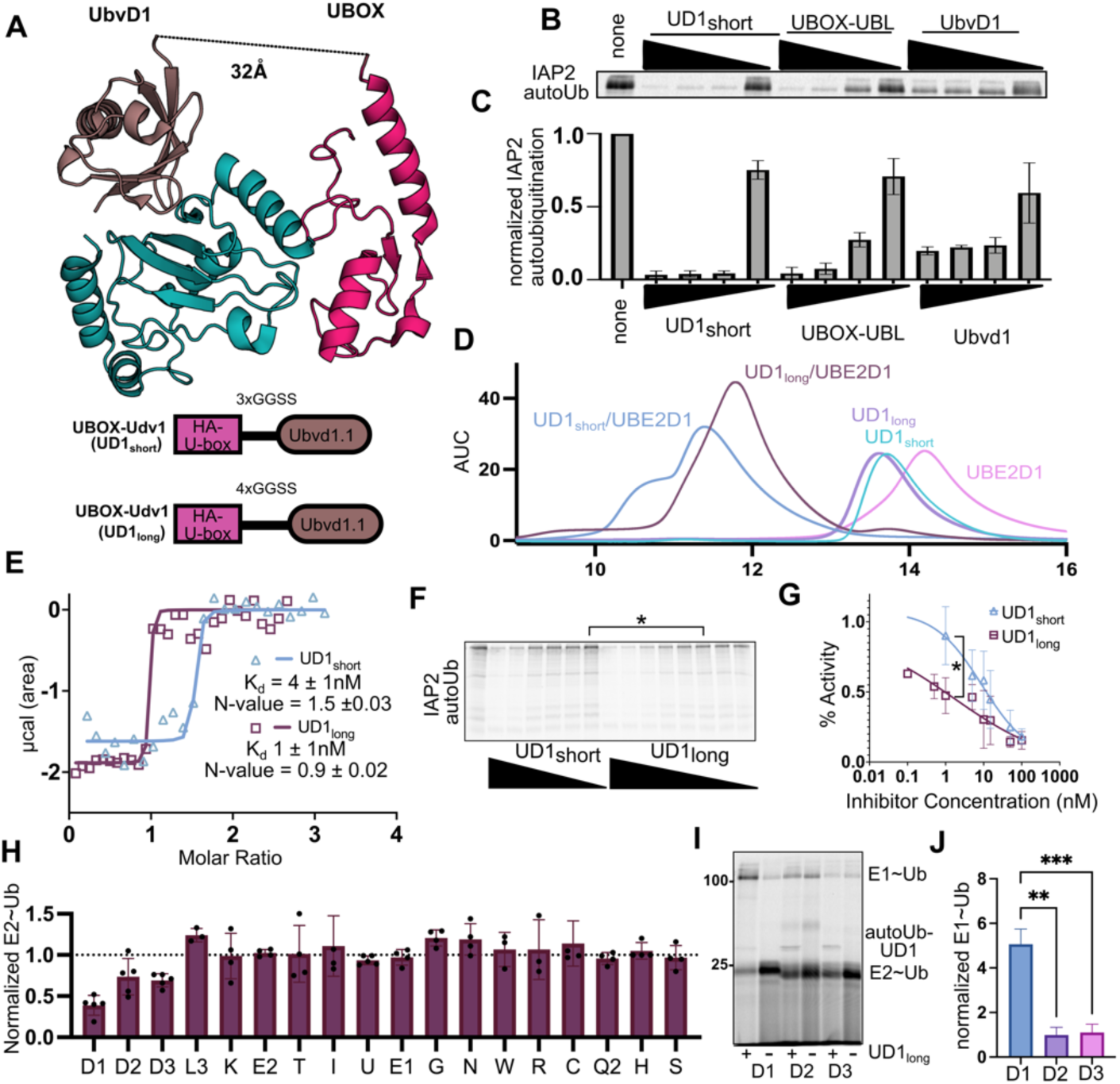
Design of high-affinity UBE2D inhibitors. *A;top*) Crystal structure of UbvD1 (PDB: 6D4P) bound to UBE2D1 with UBOX domain superimposed. *A;bottom*, Architecture of UBOX-UbvD1_short_ and UBOX-UbvD1_long_. *B)* Autoubiquitination of IAP2 in the presence of 0.1, 1, 10, and 100μM UBOX-UbvD1_short_, UBOX-UBL, or UbvD1. This assay was conducted with FITC-Ub. *C)* Quantification of the assay depicted in panel B (n=2). *D)* SEC assay showing UBOX-UbvD1_short_ (sky blue) or UBOX-UbvD1_long_ (purple) and UBE2D (pink) alone compared to the complexes (UBOX-UbvD1_short_/UBE2D1;blue or UBOX-UbvD1_long_/UBE2D1;brown). *E)* ITC binding isotherm showing the binding of UBOX-UbvD1_short_(blue) or UBOX-UbvD1_long_(brown) with UBE2D1. Heat per injection and the thermodynamics parameters are shown in Figure S4F. *F)* Autoubiquitination of IAP2 using 3nM UBE2D in the presence of decreasing concentrations of UBOX-UbvD1_short_ (100, 50, 30, 10, 5, and 1 nM) and UBOX-UbvD1_long_ (100, 50, 30, 10, 5, 1, 0.5, 0.1 nM). This assay was conducted using Cy5-Ub. *G)* Quantification of assay depicted in panel F). Statistical significance tested using a repeated-measure one-way ANOVA (*=p-value <0.05, n=3). H) E2 loading assay using UD1_long_. There is no statistically significant difference between UBE2D1, UBE2D2, and UBE2D3. UBE2D1 is statistically significant from all other E2s, while UBE2D2, and UBE2D3 are not (p-value >0.05 or greater). These assays were conducted with Cy5-Ub. *I)* E2-loading assay showing increase in E1∼Ub only in the presence of UBE2D1 and not UBE2D2 or UBE2D3. *J)* Quantification of the assay depicted in panel J. Statistical significance tested using a repeated-measure one-way ANOVA (**=p-value <.01, ***=p-value 0.001 n=5).

To further test the effectiveness of UBOX-UbvD1_short_, we determined that 3nM UBE2D1 was the lowest concentration that could support IAP2 autoubiquitination (**Figure S4B**) and used this concentration of UBE2D1 for inhibition assays. In the low E2 assay we observe robust inhibition at concentrations as low as 20nM, demonstrating the potency of the third-generation inhibitors (**Figure S4C**).

### Binding stoichiometry of the linked-domain inhibitors

Next, we wanted to test the binding stoichiometry between UBE2D1 and the inhibitors using size exclusion chromatography (SEC). We incubated RING-UBL or UBOX-UBL with UBE2D1 before running the mixture on size exclusion, which resulted in a monodispersed peak that eluted earlier than either UBE2D1 or the linked-domain inhibitor alone, and this peak has a predicted molecular weight of the 1:1 complex (**Figure S4D and S4E**). However, the chromatogram for the UBOX-UbvD1_short_/UBE2D complex yielded a heterogenous double peak and the two maxima eluted earlier than expected for a 1:1 interaction (**Figure 4D**). With this result in mind, we reexamined the co-crystal structure of UBE2D1 and UbvD1 and noticed that the C-terminus of UbvD1 is rotated compared to Ub and UHRF1 UBL. Therefore, the distance between the UBOX C-terminus and UbvD1 N-terminus is greater than the length of the 3xGGSS linker (∼2.4Å per reside), and UBOX-UbvD1_short_ cannot bind with 1:1 stoichiometry to UBE2D. We subsequently made UBOX-UbvD1_long_ with a 4xGGSS linker that could span the distance between the two domains (**Figure 4A**), and the SEC chromatogram of the UBOX-UbvD1_long_/UBE2D complex eluted as a monodisperse peak with the predicted molecular weight of 1:1 (**Figure 4D**). Using ITC, we confirmed the affinity and stoichiometry of the high-affinity binders. UBOX-UbvD1_short_ had an affinity of 4nM and had an N-value of 1.5, and UBOX-UbvD1_long_ had an affinity of ≥1nM (at or below the lower limit of detection for ITC) and an N-value of ∼1 (**Figure 4E**). These results demonstrate the importance of linker length in allowing for multivalent binding. We also directly compared both molecules in a low UBE2D concentration IAP2 autoubiquitination assay. At 1nM concentration, there is a statistically significant difference between UBOX-UbvD1_long_ and UBOX-UbvD1_short,_ and we even observed UBOX-UbvD1_long_ could still inhibit the reaction even below 1nM (**Figures 4F and G**), demonstrating the potency of the multivalent binding approach.

Finally, we examined the specificity of the UBOX-UbvD1_long_ in the E2 loading assay with 18 different E2s (**Figure 4H**). We observed selectivity for the UBE2D isoforms and no inhibition activity against other E2s. Furthermore, examining the UBE2D isoforms we observe more potent inhibition for UBE2D1 than for UBE2D2 and UBE2D3. This manifests as the persistence of E1∼Ub even after a 5-minute loading assay for UBE2D1, compared to UBE2D2 and UBE2D3 (**Figure 4I and 4J**). This specificity is anticipated since UbvD1 was designed to be selective for UBE2D1. However, when we tested UBOX-UbvD1_long_ in UHRF1 substrate inhibition assays we can observe inhibition of all UBE2D isoforms (**Figure S4H and S4I**).

Before moving into cellular assays, we wanted to ensure that the linked-domain inhibitors did not induce nonspecific ubiquitination, since we have seen that the linked-domain inhibitors can discharge the Ub conjugate. We developed a Ub promiscuity assay, where we incubated 20μM of H3 peptide with E1 and UBE2D1 at 37°C, and in the presence of UBOX-UBL or UBOX-UbvD1_long_ we can detect a small amount of non-specific peptide inhibition along with inhibitor ubiquitination (**Figure S4J**). However, these conditions are not a good surrogate for cellular ubiquitination, where many E2s are competing to be charged by E1. Therefore, we devised a multi-E2 assay to better mimic the cellular conditions. When UBE2R is added in the presence of the inhibitor and UBE2D1, the UBE2D∼Ub is significantly reduced, while UBE2R∼Ub is unaffected (**Figure S4J**). Therefore, it is unlikely that the linked-domain molecules will promote significant promiscuous ubiquitination in the cell, because other E2s will be preferentially charged instead of UBE2D. Moreover, any UBE2D that is charged will likely ubiquitinate the inhibitors, as is frequently observe in our assays.

### Expression of linked-domain inhibitors sensitize HeLa cells to cisplatin

We next set out to validate that the linked-domain inhibitors were functional in cells. We cloned the cDNAs encoding the linked-domain inhibitors into the pcDNA3.1 vector with an N-terminal FLAG tag, transfected the inhibitors into HeLa cells, and confirmed the expression using an anti-FLAG Western Blot (**Figure S5A**). We did not observe any noticeable impacts on the cellular morphology. However, when performing a careful growth assay using MTS, we discovered a slight reduction in growth rate for UBOX-UBL and UBOX-UbvD1_short_ and not for the UBOX-UBL_control_ (**UBOX-P1235T-UBL-F46V; Figure 2A**) or RING-UBL (**Figure S5B**). We probed changes to global ubiquitinome using a western blot against Ub, but we did not observe differences in the HeLa cells with UBOX-UBL or UBOX-UBL_control_ with or without MG132 treatment (**Figure S5C**). These results show that the linked-domain proteins are not blocking all cellular ubiquitination, as expected based on their selectivity.

The best-characterized effect of UBE2D inhibition/knockdown is sensitivity to chemotherapeutics^16,18–20^. Therefore, we tested the IC_50_ of HeLa cells for cisplatin, when transfected with the linked-domain inhibitors, UBOX-UBL_control_, and siRNA against UBE2D. For the linked-domain inhibitors and siRNA, we observe a reduction in the IC_50_ that scales with the affinity of the inhibitor, yet for the UBOX-UBL_control,_ the cisplatin IC_50_ was similar to HeLa cells alone (**Figures 5A, S5D, and S5E**). For example, we determined the 24-hour IC_50_ for the WT HeLa cells or those transfected with UBOX-UBL_control_ was 23-25μM (**Figures 5B, 5SD, and 5SE**), but for siRNA and UBOX-UBL, we observe a ∼3-fold reduction in IC_50,_ and for UBOX-UbvD1_short_ and UBOX-UbvD1_long_ there is a 6-fold reduction in IC_50_. Thus, the linked-domain inhibitors phenocopy UBE2D knockdown, and it appears they can outperform siRNA. Considering that the reduction of cisplatin IC_50_ scales with the K_d_ for UBE2D and the lack of chemosensitivity with UBOX-UBL_control_, these results strongly suggest that cisplatin-sensitization is the result of “on-target” UBE2D inhibition.

**Figure 5:**
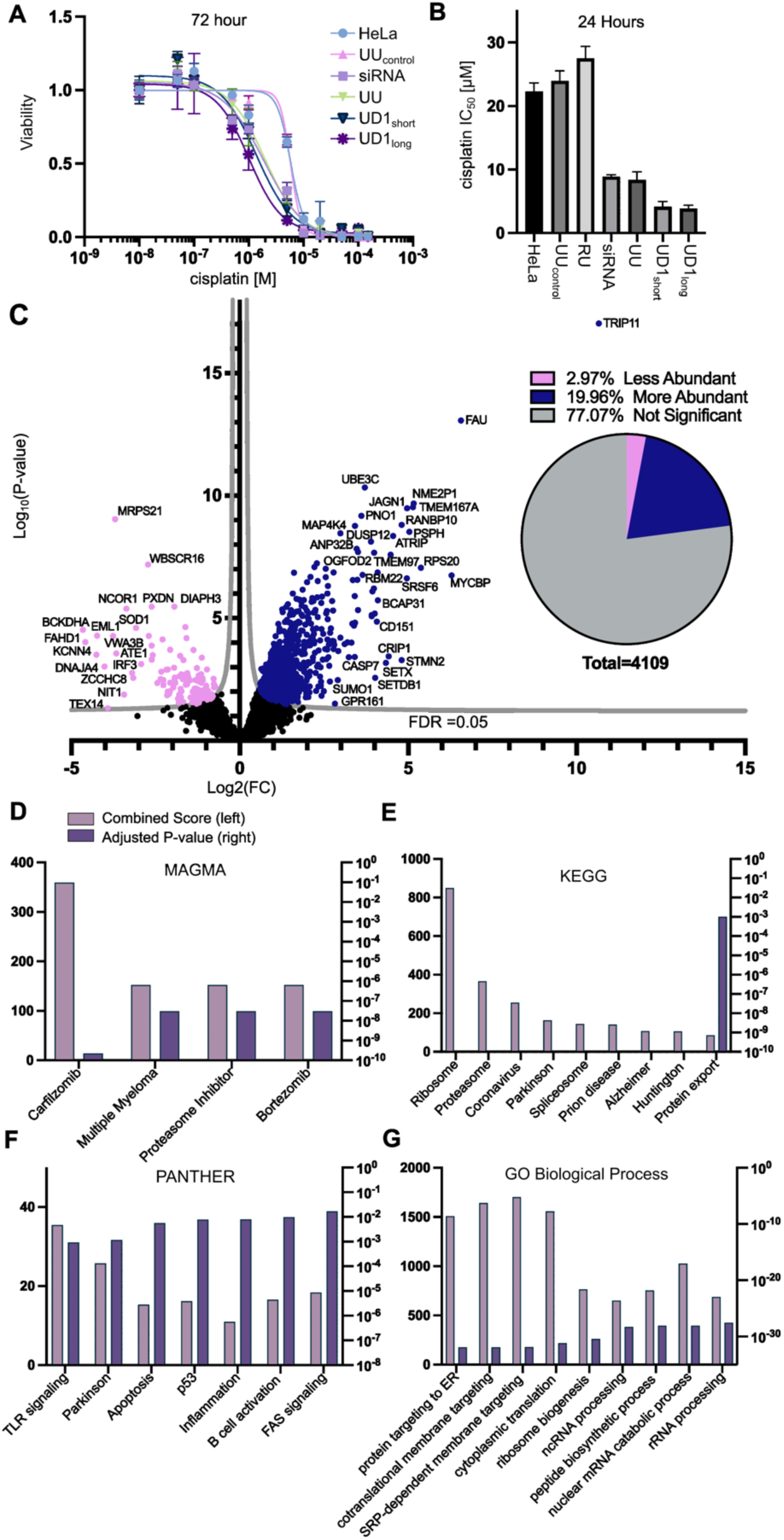
Linked-domain inhibitors rewire the proteome in HeLa cells. *A)* Viability of HeLa cells transfected with the indicated constructs and treated with indicated cisplatin concentrations for 72 hours. *B)* Bar chart of 24-hour cisplatin IC_50_ (n=6). Viability assays are shown in Figure S5D. *C; left)* Volcano plot of matched protein abundance for UBOX-UBL versus UBOX-UBL_control_. Difference is calculated as Log_2_(FC) based on label-free quantification using MaxQuant. *C; right)* Pie chart of the more abundant, less abundant, and not significantly different proteins from the shotgun proteomics. Enrichr analysis of the enriched/depleted proteins identified in shotgun proteomics tested against the indicated databases, *D)* MAGMA, *E)* KEGG, *F)* GO Biological Pathway *G)* PANTHER. The left Y-axis is combined score (pink) right Y-axis is adjusted P-value (purple).

### UBOX-UBL expression alters the HeLa cell proteome

We utilized shotgun proteomics to elucidate the impacts of UBE2D inhibition on the human proteome. Since the primary outcome of ubiquitination is protein degradation, we expected that inhibiting a central hub of the Ub cascade, like UBE2D, would increase the abundance of cellular proteins, especially those ubiquitinated by UBE2D. LC-MS/MS experiments were performed with biological and technical triplicates for each sample, providing high statistical power for our analysis. We used label-free quantification to compare the proteome of cells transfected with each inhibitor versus cells transfected with the control (UBOX-UBL_control_). In the UBOX-UBL versus UBOX-UBL_control_ comparison, we observed dramatic changes to the proteome (820 more abundant (20%) versus 122 less abundant (3%) of the identified proteins) (**Figures 5C and 5D and Supplemental Table 2**). Overall, this result is consistent with the expected function of the inhibitors, blocking ubiquitination and subsequent turnover of proteins. Remarkably, in the same experiment with RING-UBL, there were no proteins whose levels were significantly different except the two inhibitors (**Figure S5F**). This suggests that RING-UBL does not have a high enough affinity to inhibit UBE2D in cells. For UBOX-UbvD1_short/long_, the cell growth was reduced enough to complicate label-free quantitation approaches.

### UBE2D inhibition in cells mimics proteotoxic stress

To understand the biological implications of the proteomic data, we performed gene enrichment analysis of the significantly altered proteins using the Enrichr server^41,42^, which compares sets of genes against many different databases. (**Figures 5E-H and S5G-K, and Supplemental Table 3**). These results indicate that the cells are experiencing signs of reduced ubiquitin-mediated degradation and proteotoxic stress. For example, compared against the MAGMA database (Multi-marker Analysis of GenoMic Annotation)^43^, three of the four top hits were proteasome inhibitor treatments (Carfilzomib/proteasome inhibitor/Bortezomib) (**Figure 5E**) and in the KEGG^44^ and PANTHER databases, the top hits include protein aggregation diseases, like Parkinson’s, Prion, Alzheimer’s, and Huntington’s. These results further suggest that the linked-domain proteins are partially inhibiting Ub-mediated proteasomal degradation (**Figure 5F and G**).

We also observe changes to many complexes and pathways required for proteostasis, such as the ribosome, proteasome, protein transport, and somewhat surprisingly, many nuclear RNA processes, like splicing, RNA processing, and nuclear mRNA processing, suggesting potential nuclear-specific functions of UBE2D. Importantly, a recent study using RNAi to knockdown E1 and E2s showed that upon reducing UBE2D levels, there is an increase in peroxisomal proteins^45^. We also observe an increase PEX3, which was the most increased PEX protein in Hunt et al. and was confirmed by Western Blot. Additionally, we observe PEX16, as well as other peroxisomal proteins SLC25A17, PRDX1, IDH2, FAR1, and SOD2. Therefore, inhibition of UBE2D appears to activate an adaptive stress response to decreased ubiquitination. The linked-domain inhibitors described here will be a useful tool for future targeted studies to dissect the basis for this response and other UBE2D-specific functions in the cell.

## Discussion

In this study, we engineer a new class of high-affinity, protein-based inhibitors that are highly selective for the UBE2D family of E2-conjugating enzymes. The linked-domain inhibitors have affinities ranging from 10^-6^M to <10^-9^M and are the most selective and highest affinity (1,000-10,000-fold higher affinity than small molecules) inhibitors of UBE2D described to date. These tools can be spatial-temporally regulated and will allow researchers to ask previously impossible questions about the cellular role of UBE2D. The potency of these molecules is due in part to directed evolution, but also because of the multivalent interactions with E2s that were drawn from native interaction with E3s.

The linked-domain inhibitors are active in HeLa cells and increase the abundance of ∼20% of the identified proteins, consistent with a reduction in Ub-mediated protein degradation. The impacted biological pathways span most aspects of proteostasis (transcription, translation, protein localization, and protein degradation). The genetic profiles of the cells, compare to cells that have reduced Ub-mediated degradation or experiencing proteotoxic stress. When our results are viewed within the context of the broader literature, it suggests that UBE2D activity is an important hub in this response, and decreasing its activity leads to an adaptive response and decreased resiliency to stress^45,46^. This is supported by the 6-fold reduction in HeLa-cisplatin IC_50_ when treated with the most potent UBE2D inhibitor (**Figure 5B**). The gene enrichment analysis suggests a mechanism for the observed reduced fitness because apoptosis/ferroptosis was identified as hits in several databases (**KEGG; Figure 5E, Panther; Figure 5H, MSigDB; Figure S5G, and BioCarta; Figure S5K**) ^47,48^.

Indeed, we observe increases in many proteins involved in apoptosis such as the executioner Caspase 7, the proapoptotic mitochondrial protein SMAC/DIABLO^49^, the inositol 1,4,5-trisphosphate receptors ITPR1 and ITPR3 that trigger calcium release from the endoplasmic reticulum^50^, the lysosomal protease cathepsin Z, the UBL containing protein DFFA, which fragments DNA during apoptosis, HMG1/2, and the UBL MAP1LC3B, which has been shown to induce apoptosis upon proteasomal inhibition^51^. There are also changes to FAS-mediated apoptosis, such as an increase in FAF1, a potentiator of apoptosis, and decreased levels of DAXX, a protein that inhibits apoptosis^52^. Interestingly, decreased DAXX was also observed in Hunt et al. when treating the cells with UBE2D RNAi, suggesting that this is a signaling event that occurs in response to UBE2D inhibition. Additionally, we observe proteomic signatures of mitochondrial stress, such as inner membrane proteins involved in oxidative phosphorylation (cytochrome C oxidase and reductase, NADH dehydrogenase, succinate dehydrogenase, and the F-type ATPase), and mitochondrial ribosomal subunits.

We also detect changes to protein levels involved in NF-kB signaling/inflammation^53^ (**Figure 5H; Supplemental Table 3**). We observed increased IKK-α and RelA (p65), one of the NF-kB transcription factor subunits, and decreased catalytically active IKK-β subunit, a primary driver of prosurvival inflammation. These changes appear to indicate a reduction of prosurvival NF-kB signaling and is consistent with the anti-inflammatory activity observed for chemical inhibitors of UBE2D^17^. The combination of proapoptotic and anti-inflammatory effects from inhibiting UBE2D may contribute to decreased sensitivity to cisplatin and warrants further evaluation of UBE2D therapeutically.

One of the outstanding questions in Ub biology remains how networks of Ub machineries work together to regulate the proteome in cells. These questions are difficult to address because of a lack of specific tools, and we anticipate that our linked-domain tools will be useful to disentangle E2, E3, and deubiquitinase relationships. Indeed, we observed changes to 3 E2s, 9 E3s, and 3 DUBs in response to the linked-domain inhibitor (**Supplemental Table 4**). Interestingly, the E2s are UBE2Z and UFC1, noncanonical E2s that can work with other conjugatable UBLs, and we also observe increased levels of the UBLs FAU (FUBI), MAP1LC3B, which is involved in autophagy, and SUMO1. It is attractive to speculate not only about reprogramming of the ubiquitinome, but also reprogramming of the UBLome in response to reduced ubiquitination. We envision future work to make a suite of linked-domain E2 inhibitors that will enable pinpointing the roles of the Ub-conjugating enzymes in cells.

## Acknowledgements

Our work is supported by start-up funds from UoP (JSH), Instrumentation grants (MRI-1828179 and DBI-1531417), Stauffer Research Undergraduate research grants (A.C, K.R.H, G.L), NIH R35GM128855 and UCRF (N.G.B), NIH T32GM008570 (D.L.B), and R01GM141409 (G.K and S.H.)

## MATERIALS AND METHODS

### Cloning

All linked-domain inhibitor genes were ordered from TWIST biosciences and cloned into either a modified version of the pQE-80L vector which contains his-MBP with a TEV-cleavage site (RING-UBL, UBOX-UBL, UBOX-UbvD1_short_, UbvD1, UHRF1-UBL, UHRF1-RING, high-affinity UBOX, IAP2) or the pET44 vector with the NusA tag removed (UBOX-UbvD1_long_) and a TEV cleavable N-terminal 10x Histidine tag. Mammalian expression constructs were cloned into pcDNA 3.1 with an N-terminal FLAG tag. The UBL sequence consisted of UHRF1_1-76_, the RING sequence was UHRF1_675-793_, the UBOX sequence was UBE4B_1221-1302 L1236I/I1252V_. Inhibitor protein sequences are found in Supplemental Table 5. All E2s were also expressed as MBP fusions in the afore mentioned vector, except for UBE2D1 and UBE2E1, which had N-terminal polyhistidine tag.

### Protein Expression

Bacterial expression plasmids were transformed into BL21-CodonPlus Competent Cells (Agilent) using heat shock. Starter cultures were then grown overnight (ON), and 1L cultures were inoculated with 1:100 dilution and grown for at 37°C until they reached O.D. 600 value of 0.4-0.6. IPTG (Gold Biosciences) was added at a 1mM and cells were induced O/N at 16°C and 200 rpm. Cells were harvested by centrifugation and pellets were frozen were collected for further purification.

### Protein Purification

The standard purification for his-MBP and his tagged proteins is to resuspend the cell pellet in lysis buffer (50 mM TRIS-HCl, pH 8.0, 300mM NaCl) with PMSF (500μM) and bestatin (10μM). The lysate was sonicated on ice and clarified by centrifugation. All lysates were run over an Ni-NTA resin (Gold Biosciences), washed using 50-100 mL of 50mM TRIS-HCl, pH 8, 1M NaCl, 5 mM imidazole, and eluted using 10-20 mL of elution buffer (50 mM TRIS-HCl, pH 8.0, 100mM NaCl, 250mM imidazole). The protein was dialyzed overnight at 4°C into 50mM HEPES, 100mM NaCl, 1mM DTT, pH 7.4. his-MBP tag constructs were cleaved using TEV protease purified in house. The cleaved His-MBP tag was typically removed using the NiNTA resin or using ion exchange column (Buffer A: 50mM Hepes, 100mM NaCl pH 7.4, Buffer B 50mM Hepes, 500mM NaCl). Fractions were pooled, concentrated using centrifugal filter units, and then run over the size exclusion column (Sephacryl S200; Cytivia) (Buffer: 50 mM HEPES pH 7, 100 mM NaCl, 1 mM DTT). Proteins were concentrated and frozen for future assays. Other proteins used in this study were purified according to previous reported methods. Ub-G76C was produced for fluorescent labeling and was purified as a GST fusion, cleaved with TEV, then labeled using bifunctional maleimide fused fluorophore, either FITC-maleamide or Cy5-maleamide (Cayman Chemical). UBE2D1 and UBE2E1 have a N-terminal poly-his tag and was purified using Ni-NTA by size-exclusion and UHRF1 was purified using the standard his-MBP vector. Cul3/Rbx1 were expressed using the “split and co-express” system in E. coli, and purified sequentially using Ni-NTA followed by GST. The protein was cleaved off the resin using TEV and then run over (Sephacryl S200; Cytiva) as previously reported^54^. APC/C was expressed in insect cells and purified to high homogeneity by using a C-terminal twin-Strep tag on APC4, then anion exchange chromatography and gel filtration as previously described^38^. βTrCP-Skp1 was expressed in insect cells and purified using glutathione resin, cleaved with thrombin, then further purified using anion exchange, and size exclusion as previously reported^55^. The Cul1-Rbx1 was purified using the split and co-express system described above for Cul3-Rbx1. Neddylation and isolation of the of the modified Cul1-Rbx1 was done according as previously reported^56^. UBE2D2 and UBE2R used in these assays were purified as GST fusions and cleaved from the resin. His-SUMO-UBOX, UBE2E1, and UBE2D1 were purified according to the standard His protocol^27^. E2s used in the panel assays were expressed as His-MBP fusions and purified as described above and the cleaved E2s were isolated by passing them over the Ni-NTA column.

### Ubiquitination Assays

All cell free ubiquitination assays were carried out in a total reaction volume of 20µL. The reaction consists of 50mM HEPES pH 7.4, 2.5 mM MgCl2 2.5 mM DTT, 100mM NaCl, 10mM ATP, 5-20µM of the indicated Ub (FLAG, Fluorescent), E1 activating enzyme 50-100nM, and 675nM of Ube2D1, unless indicated otherwise. Low concentration UBE2D1 assays were performed by using 3nM UBE2D1. Reactions were quenched with SDS-PAGE sample loading buffer devoid of reducing agents. 7µl of samples were loaded onto 12-15% SDS-PAGE gels and subsequently imaged using the STORM 860 Molecular Imager (Molecular Devices, San Jose, CA, USA) and gel fluorescent bands were quantified using densitometric analysis through ImageQuant Software (v5.2)/ Gelanalyzer (V19.1). All bands were background corrected and were subsequently normalized to the positive control reaction to plot relative activity. E3s (UHRF1, IAP2, Cul3) were added at 1μM. For substrate assays with UHRF1, hemi-methylated DNA (IDT-DNA) was added at 3μM and H3_(1-25)_ peptide (BioMatik) was used at 10μM according to previous studies^28^. APC/C substrate ubiquitination assays were conducted with 30nM APC, 100nM E1, 2µM E2, 500nM CDH1, 200nM Ub-Cyclin B (produced as previously described), 100µM Ub, 5mM ATP, and BSA and quenched at 8 minutes similar to other reported assays^36^. SCF substrate reactions were conducted using the following conditions 500 nM E1, 100nM neddylated SCF (β-TRCP), 5µM β-catenin peptide, 60µM Ub and, 2µM of the respective E2. Reactions with UBE2D2 were incubated for 5 minutes while reactions with UBE2R were incubated for 15 minutes because of slower kinetics^55^. UBE2D-charging assay contained 100nM E1, 2µM UBE2D2 and 4µM Ub and time points were taken at 5, 15, 45 and 90 seconds. E1 competition assays had increasing E1 from 50nM to 200nM in the presence of 1µM UBOX-UBL with no E3 ligase present. The reaction was quenched at 90 seconds. For UHRF1 competition assays, UHRF1 concentration was varied from 0.7, 2, 4µM in the presence or absence of a 15µM RING-UBL. The reaction mixture contained 19µM H31-25 peptide, 1µM HeDNA, and 675nM UBE2D1. Reactions were run for 20 min at RT. The double E2 loading assay were conducted at 37°C and contained UBE2D1 (5µM) and UBE2R1 (5µM), +/- 19µM H3 peptide, and +/- 10µM inhibitor. The reactions were initiated by adding the reaction mixture and the assays were run for five minutes and quenched using non-reducing SDS-page gel. All statistical tests were carried out in PRISM.

### Ub Discharge Assays

Thioester discharge assays were conducted at room temperature with 1µM E1, 2µM UBE2D2, and 2µM fluorescent Ub, K_0_ (ubiquitin with all lysines mutated to arginine) and ran for 30 minutes before adding 50mM EDTA to quench the reaction. Then 23µM of UBOX-UBL, RING-UBL, UBL, UHRF1-RING, or HA-UBOX were added with or without 20mM lysine or UbΔGG (1-74). Oxyester discharge assays were conducted as previously described^25^. Briefly, the oxyester-UBE2D is purified using S75 (superdex increase; Cytivia), then added at 15 µM to the solution containing the indicated amount of each inhibitor and 20mM lysine and quenched at the indicated time points with reducing dye. The results were visualized using Coomassie staining. Importantly this assay does not use EDTA, so this shows the lack of activity from the UHRF1-RING is not due to a loss of zinc. The purified oxyester-UBE2D was a generous gift from Rachel Klevit’s lab.

### E2 Panel Loading Assay

Eighteen E2s were added at 5µM with or without 10µM inhibitor. The reaction mix containing the above-mentioned components, except E2. The reactions were initiated by adding the E2s and allowed to proceed for 5 minutes. The assays were quenched with non-reducing loading dye and immediately ran on an SDS-page gel and visualized using the Storm scanner to visualize the fluorescent Ub.

### Isothermal Calorimetry

UBE2D1 and inhibitor were dialyzed into 25mM HEPES pH 7.0, 100mM NaCl, 1mM TCEP. ITC experiments were performed using the Affinity ITC LV (Waters, TAInstruments). 1.5µL injections of linked-domain inhibitors were injected into an isothermal cell containing UBE2D1. Experiments were performed at 25°C. The delay between each injection was 300 seconds. A heat-burst curve was generated (micro calories/second vs. seconds) for each injection and the area under the curve was calculated for each injection using NanoAnalyzer software (version 3.8.0) to determine the heat (kJ/mol) associated with each injection. The last 5 injections were used to determine a blank constant that was used to adjust the raw measurements. The dissociation constant was also determined using NanoAnalyzer Software (version 3.8.0) after fitting the adjusted measurements to an independent model.

### Yeast Two Hybrid

Linked-domain inhibitors were cloned into the BamH1 and EcoR1 sites of the pGTKT7 (Takara) vector, which fuses the GAL4-DNA binding domain to the N-terminus of gene. The E2 GAL4AD fusion vectors were a gift from Rachel Klevit’s lab^39^. Screening was performed by stepwise transformation of each vector into Y2H Gold yeast (Takara). Co-transfected yeasts were grown in liquid culture starting from glycerol stocks. After overnight growth in YPD, cultures were transferred into synthetic media (-His/-Trp/-Leu) with Aerobasidin A. Growth was assessed by measuring the optical density at 600nm between 5-13 days.

### Size Exclusion

Proteins were incubated 1:1 at 20μM concentration for 10 minutes before being run over the Superdex 75 10/300 GL column. Molecular weight curve was generated using proteins commonly produced in lab of varying molecular weights.

### Cell culture

HeLa cells were obtained from Dr. Willian Chan’s lab (School of Pharmacy, University of the Pacific) and cultured in EMEM media supplemented with 10% fetal bovine serum (FBS) and 1% penicillin-streptomycin solution (10,000 U/mL penicillin, 10,000 μg/mL streptomycin). Cell cultures were incubated at 37°C in a humidified atmosphere of 5% CO2 and 95% air.

### Cytotoxicity assay

The cytotoxicity of cisplatin was measured in untransfected HeLa cells and HeLa cells transfected with the control expression vector (UC), the four inhibitors expression vectors (RING-UBL, UBOX-UBL, UBOX-Ubv1_short_ and UBOX-Ubv1_long_) and the UBE2D siRNA (Santa Cruz) using MTS (Promega) tetrazolium assay. After three passages, cells were seeded onto 6-well plates with seeding number of 1.5×10^5^ cells/well and were allowed to grow for 24 hours before the transfection. Right before transfection, old medium was extracted and 4 mL of complete EMEM medium was added to each well. DNA to be transfected to the cells in each well was dissolved in 400μL EMEM medium with a concentration of 10ng/μL and incubated for 5 minutes. Then, 6μL TurboFect (ThermoFisher) was added into the above mixture and incubated for 18 minutes. Afterwards the DNA mixture was pipetted on the cells. After 24 hours, cells were harvested by trypsinization, counted and calculated by EVETM cell counter from NanoEn Tek using trypan blue dye exclusion. Then cells were plated at 4000 cells/well by adding 100μL of 4×10^4^ cells/mL suspension solution into each well of the 96-well culture plate. Cells were allowed to grow for 24 hours before cisplatin treatment. The stock solution of cisplatin was freshly prepared in autoclaved 0.9% NaCl. Then the mixture of complete EMEM medium and cisplatin solution were prepared to achieve each concentration of cisplatin in a wide range from 10nM to 150μM. Right before the treatment, old medium was extracted and replaced with 110μL cisplatin mixture to obtain a dose-response curve for 24, 48 and 72 hours. 20μL MTS reagent was added to each well 3 hours before reaching each time point. Then the absorbance of the MTS-formazan product at 490 nm was measured with the microplate reader INFINITE M PLEX (TECAN).

### Sample preparation for Mass Spectrometry

48 hours post transfection Hela Cells were scraped and washed with a PBS buffer and pelted down at 4°C. The cells were then lysed using 200μL of lysis buffer (50mM HEPES, 150mM KCl, 10% glycerol, 0.5% NP-40, 1mM EGTA, 1mM MgCl2). Protease inhibitors were added (PMSF 500μM and Bestatin 20μM) and 20 G X ½ syringe pass was done five times at 4°C to lyse the cells. The lysed cells were then centrifuged at 17,000 G for 30 min.

### Reduction and Alkylation

The supernatant was collected and reduced by adding 5mM of DTT and incubated at 37 for 1 hour. The samples were further alkylated by addition of 50 mM iodoacetamide and were incubated at room temperature in the dark for 20 minutes. 60μL of lysis buffer was added post the incubation and samples were vortexed then the reaction was quenched using DTT.

### Sample treatment for MS

Samples were precipitated by addition of 400 μL of MS grade chilled methanol. Rigorous mixing was achieved by vortexing. 100μL of chilled chloroform was further added to the samples followed by addition of 300μL of chilled Milli-Q water. The samples were vortexed and then centrifuged at 17,000 X G at 4 The aqueous layer was removed without disturbing the interphase containing the protein. 300μL of MS grade methanol was further added and samples were vortexed and centrifuged at 17,000 X G at 4°C for 5 minutes. The supernatant was discarded, and the pellet was dried down using Speed Vac.

### Trypsin Digestion

The pellet was resuspended in 50μL of 50mM ammonium bicarbonate and 10μL thawed trypsin (Promega) was added. The sample were incubated overnight at 37°C. Additionally 5μL of Trypsin was added the next day followed by 4 hours of incubation at 37°C the next day. The samples were cleaned up using C-18 Spin columns (Pierce) and diluted 1:10 times in water with HPLC .1% Formic Acid for LC-MS/MS.

### Instrument Parameters

Mass spectrometry analyses were performed using an Orbitrap Fusion™ Tribrid™ mass spectrometer equipped with an EASY-Spray™ ion source (Thermo Fisher Scientific) operated in a data-dependent acquisition (DDA) manner by Xcalibur 4.0 software (Thermo Fisher Scientific). Samples were loaded onto the column for 10 min at 0.300 μL/min using a previously described gradient ^57^. Wash runs were conducted in between each sample and each sample was run in biological triplicate with technical triplicates. MS1s were collected using the orbitrap in positive mode using a resolution of 120000, a scan range of 400-1600, ACG target of 1.0×10^6^, and a maximum injection time of 50 seconds. MS2s were collected in using the ion trap with Turbo scan rate, ACG target of 1.0 x10^4^ and maximum injection time of 35ms.

### Identification, quantification and statistical analysis

MS raw files were analyzed using MaxQuant software 2.0.3.1 with the Andromeda search engine. Searches were performed against the Uniprot database for Homo sapiens (UP000005640, May 2022). UBOX-UBL, RING-UBL and UBOX-UBL_control_ protein fasta sequences were initially also added and searched against. Replicates were grouped and LFQ quantitation was separated in parameter groups. For identification, carbamidomethylation was set as a fixed modification and N-terminal acetylation and methionine oxidation as variable modifications. Statistical analysis of the MaxQuant result table proteinGroups.txt. was done with Perseus 1.6.14.0. Potential contaminants, reverse peptides and peptides only identified by site were removed. Raw intensities differences were Log2-transformed. Rows were then divided into two groups: inhibitor (UBOX-UBL or RING-UBL) transfected samples or samples transfected with the negative control (UC). At least 3 valid values in at least one group for each row was required and filtered on. Missing values were replaced from the normal distribution separately for each column with a down shift of 1.8. Two-sided t-tests were performed to obtain FDR corrected *p*-values (FDR=0.05) using the Permutation-based FDR function. The mass spectrometry proteomics data have been deposited to the ProteomeXchange Consortium via the PRIDE partner repository with the dataset identifier PXD040264.

### Enrichr analysis

Proteins that had statistically significant FDR corrected *p*-values in the UBOX-UBL sample versus UBOX-UBL_control_ were searched against a variety of databases using the Enrichr website. Top hits were plotted and all results were included in the supplemental table 4.

---

**Supplemental Figure 1:**
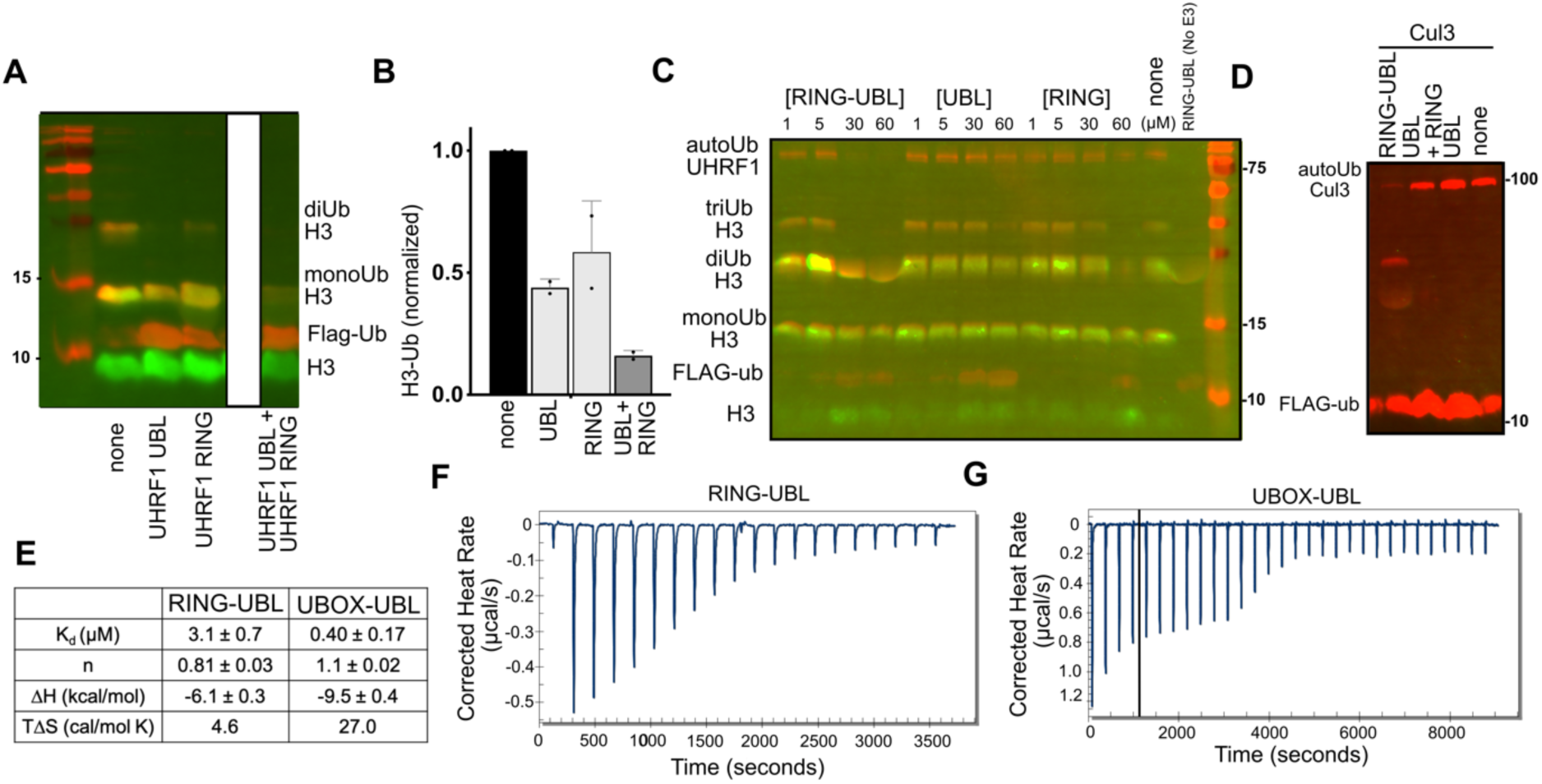
*A)* Representative blot of UHRF1 histone ubiquitination assay incubated with UHRF1 UBL (50μM), UHRF1 RING (75μM), and UHRF1 UBL (50μM) and UHRF1 RING (75μM) together. Ub is visualized using anti-FLAG Western Blot and peptide is visualized using strepatavadin-488. *B)* Quantification of the assay depicted in Figure S1A (n=2). *C)* UHRF1 ubiquitination assay comparing RING-UBL to the UBL and RING domain alone. *D)* Cul3 autoubiquitination assay comparing RING-UBL to the UBL and RING+UBL. *E)* Thermodynamic parameters from fitting ITC data for RING-UBL and UBOX-UBL. Heat per injection plots for RING-UBL *(F)* and UBOX-UBL *(G)*. To fit the curve shown in panel G, we excluded the first four points due to spurious heat release not from UBOX-UBL/UBE2D binding that was not present in other ITC runs.

**Supplemental Figure 2:**
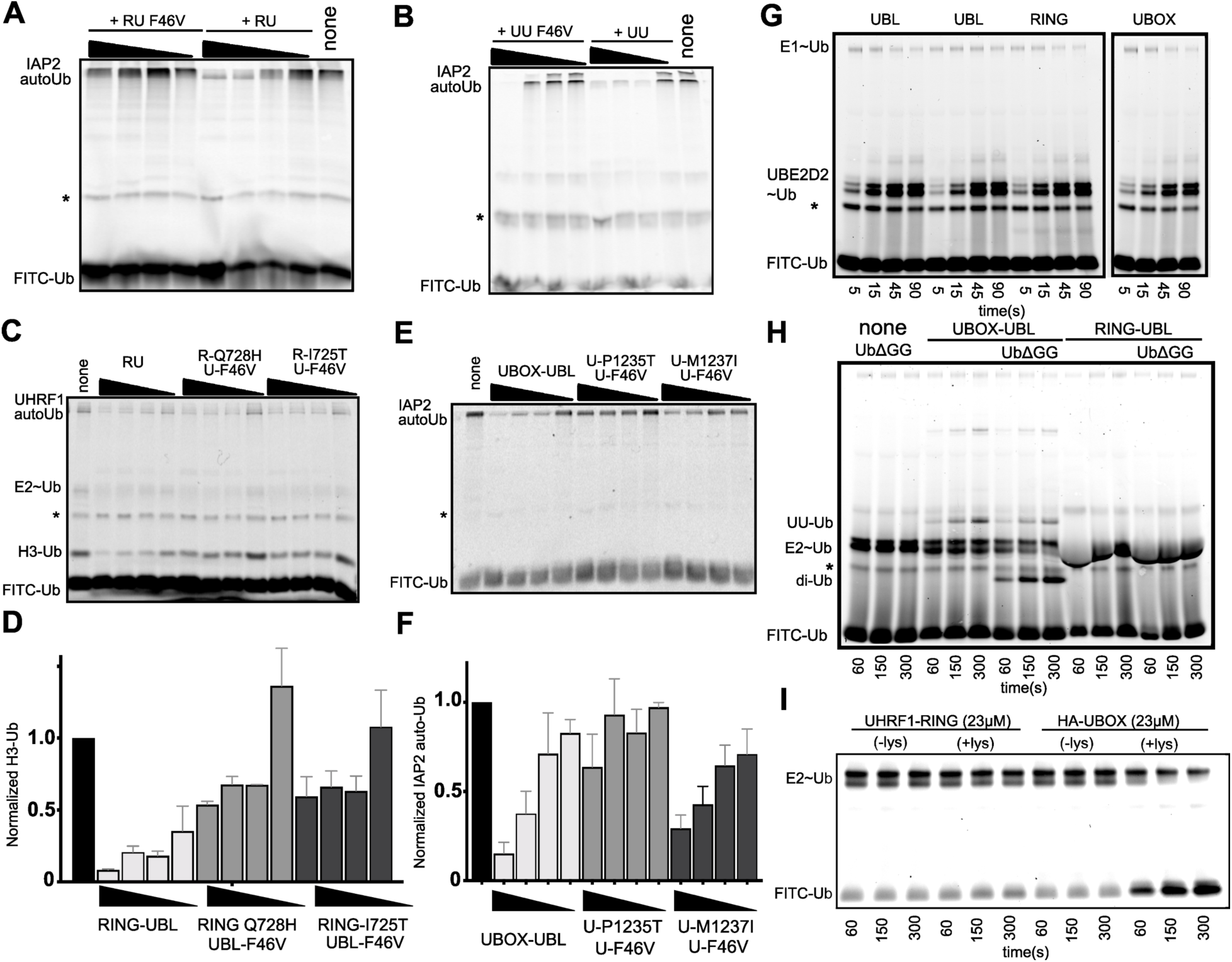
IAP2 autoubiquitination assay with the F46V mutation in RING-UBL *(A)* or UBOX-UBL *(B)*. C) UHRF1 ubiquitination assays with the corresponding RING-UBL mutations. *D)* Quantification of C (n=3). *E)* IAP2 autoubiquitination assays with the corresponding UBOX-UBL mutations. *F)* Quantification of E (n=3). In assays A-F inhibitors were loaded at 100, 50, 10, and 1μM. *G)* E1 loading assay for the indicated constructs at 23μM. Two different purifications of UBL were used in this experiment and this assay was run in parallel with the assays in Figure 2C. *H)* Ub discharge assays from UBE2D in the presence of the 23μM indicated proteins. *I)* Lysine discharge assay for the indicated proteins. All Ub assays in the panel contained FITC-Ub and the relevant bands are labeled. The * corresponds to a background band commonly observed in fluorescent Ub preps.

**Supplemental Figure 3:**
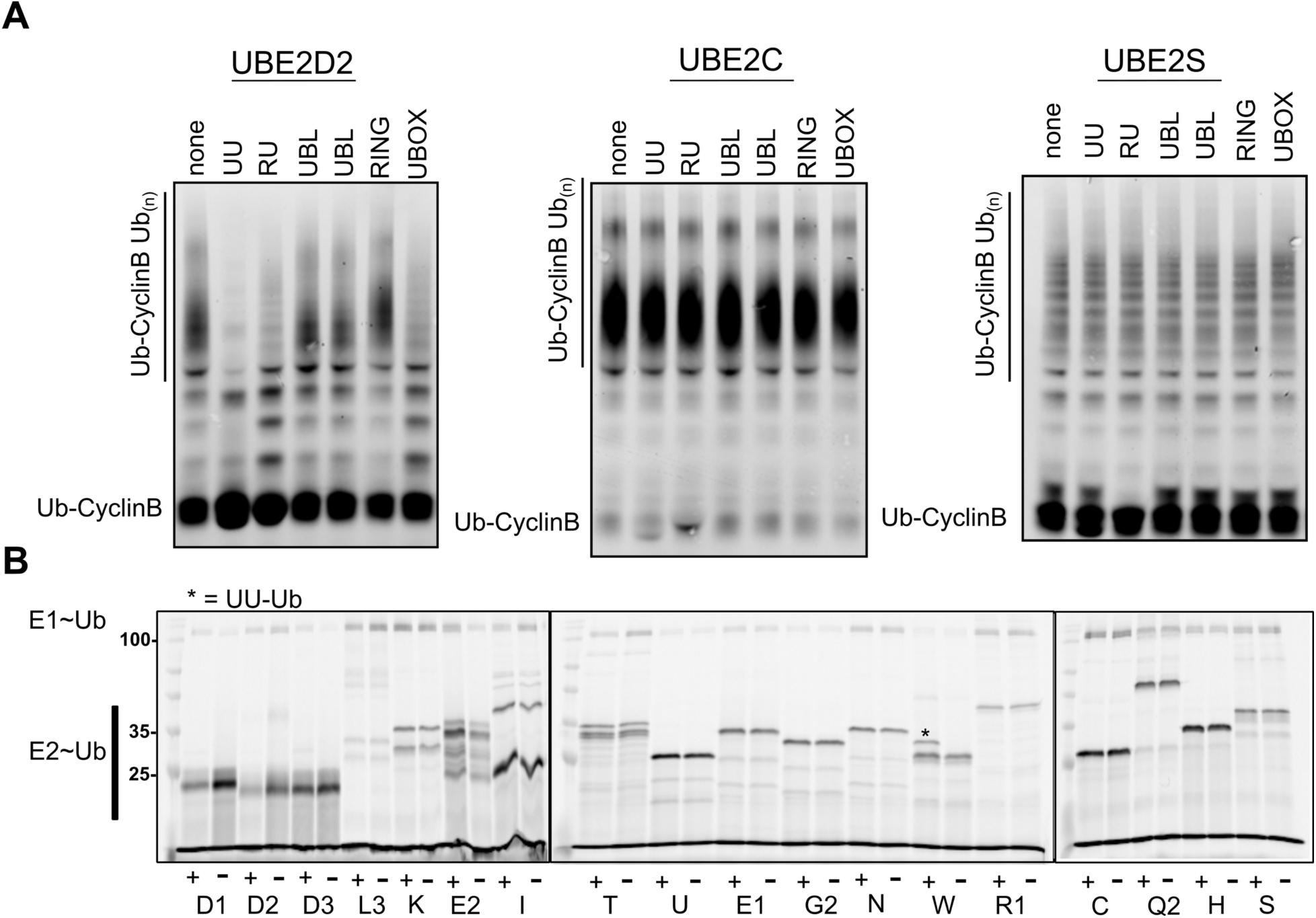
*A)* APC ubiquitination assays with the indicated E2s in the presence of the indicated proteins at 23μM and using 2μM of the indicated E2. Reactions were quenched using reducing SDS-page loading buffer at 12 minutes and fluorescent Ub-CyclinB was used to monitor the reaction. The upper polyubiquitin band was quantified and shown in Figure 3D. *B)* Representative E2 loading assay with and without UBOX-UBL and 5μM of the indicated E2. Reactions were quenched with nonreducing SDS-page loading gel at 5 minutes.

**Supplemental Figure 4:**
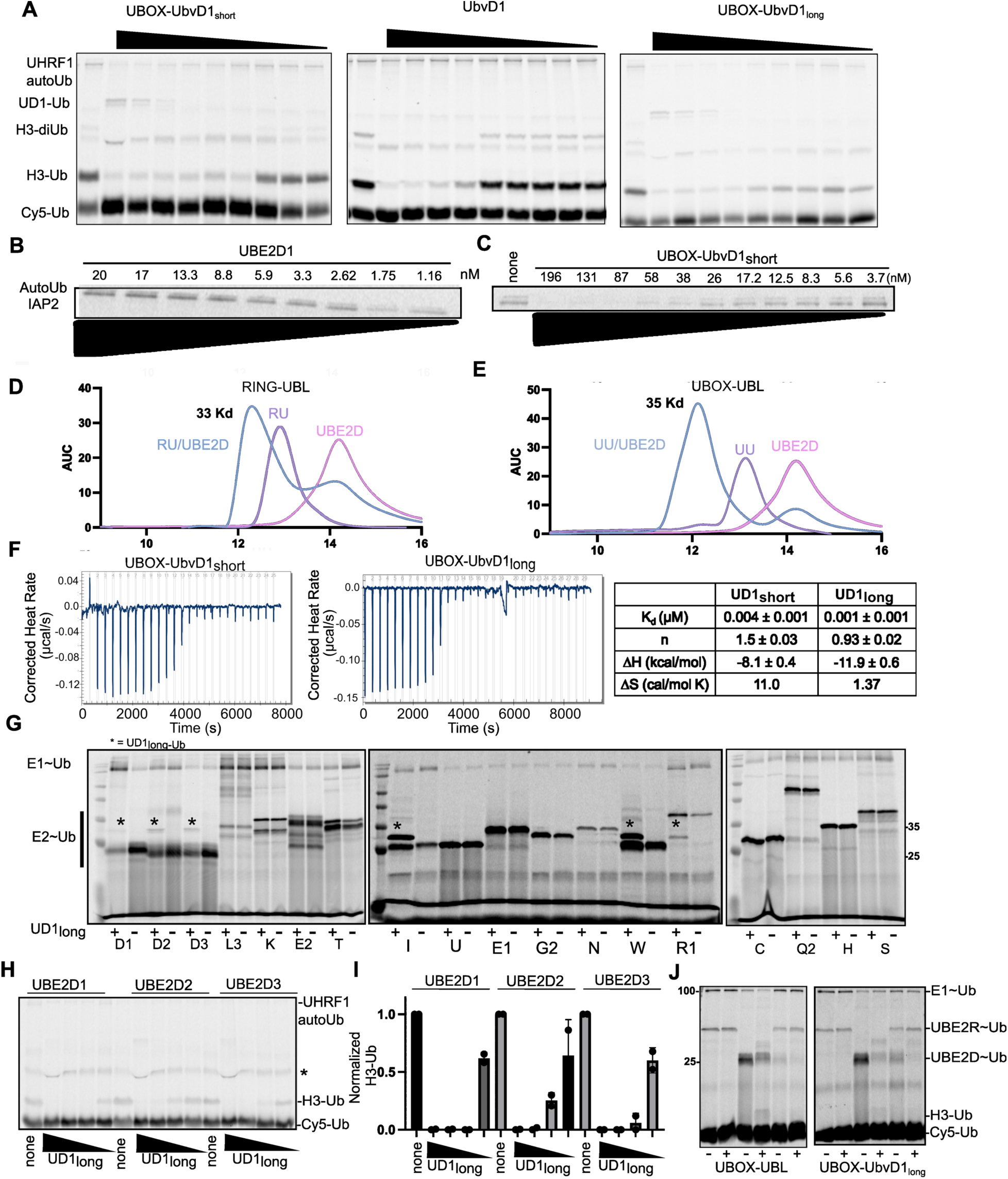
A) UHRF1 ubiquitination assays in the presence of *A)* UBOX-UbvD1_short_, UbvD1, or UBOX-UbvD1_long_. at 45, 16, 4.0, 1.0, 0.24, 0.13, 0.09, 0.06, 0.04 μM. *B)* IAP2 autoubiquitination assay with the indicated concentrations of UBE2D1. *C)* Low concentration UBE2D (3nM) IAP2 autoubiquitination assay with the indicated concentrations of UBOX-UbvD1_short_. Size exclusion chromatograms for *D)* RING-UBL or *E)* UBOX-UBL alone or in complex with UBE2D1. *F; left)* Isothermal Titration Calorimetry heat per injection plot for UBOX-UbvD1_short_ and UBOX-UbvD1_long_. *F;right)* Table of the thermodynamic parameters from the ITC fitting. *G)* Representative E2 loading assay using UD1_long_ with the indicated E2s. Quantification of the E2∼Ub is depicted in Figure 4H. The UBE2D1, 2, and 3 lanes are also shown in Figure 4I. *H)* UHRF1 ubiquitination assay using either UBE2D1, 2, and 3, in the presence of UD1_long_ 100, 10, 1, 0.1 μM. *I)* Quantification of the H3-Ub band from the assay shown in panel H. *J)* Double E2 promiscuity assay using either UBE2R, UBE2D1, or UBE2R and UBE2D1 together with and without 10μM of the indicated linked-domain inhibitors in the presence of 19μM H3 peptide. These assays were run at 37°C.

**Supplemental Figure 5:**
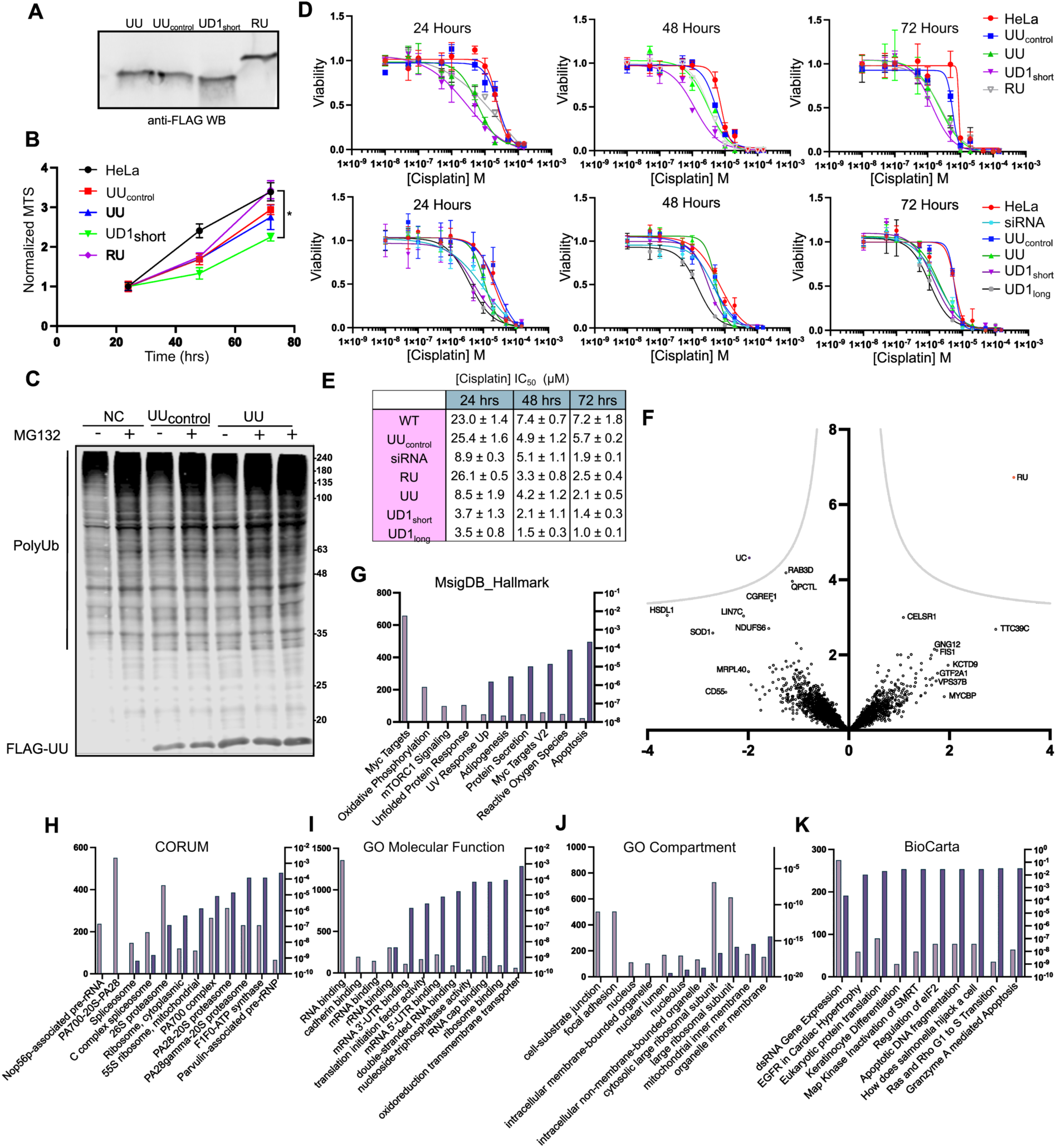
*A)* FLAG western blot to visualize the inhibitors transfected into HeLa cells. *B)* Growth assay for HeLa cells transfected with the indicated inhibitors. Growth is normalized to the 24-hour MTS value for each sample. *C)* Anti-Ub western blot of HeLa cells treated with UBOX-UBL and UBOX-UBL_control_ in the absence and presence of MG132. *D)* Cisplatin titrations for HeLa cells transfected with the indicated vectors at the indicated time points. Viability is measured using the MTS assay. 72-hour time point (bottom) is also shown in Figure 5A. *E)* Cell viability for HeLa cells incubated with cisplatin at the indicated time points transfected with the indicated inhibitors. *F)* Volcano plot of matched proteins in RING-UBL versus UBOX-UBL_control_. Enrichr analysis of the more/less abundant proteins from the UBOX-UBL versus UBOX-UBL_control_ shotgun proteomics experiments shown in Figure 5: *G)* MsigDB, *H)* CORUM, *I)* GO molecular function, *J)* GO compartment, *K)* BioCarta.

